# Long-term liana infestation leaves architectural legacies in host trees

**DOI:** 10.64898/2026.09.18.752529

**Authors:** Rosie Young, Helene C. Muller-Landau, Félicien Meunier, Hans Verbeeck, Sruthi M. Krishna Moorthy, Roberto Salguero-Gómez

## Abstract

Lianas are an integral component of tropical forests, and their increasing abundance may alter forest structure and function. Whether infested host trees are shorter and smaller-crowned because lianas suppress them, or because lianas preferentially colonise already small trees, has remained unresolved. We used a longitudinal dataset of individual liana infestation histories spanning fourteen years (2011–2025), combined with terrestrial laser scanning (TLS)-derived structural metrics for 251 trees across 67 species on Barro Colorado Island, Panama. Bayesian analyses showed that persistently infested trees were shorter (−11.0%), with smaller crown projected area (−22.3%) and crown volume (−25.9%) than never-infested trees; the 95% credible intervals for these effects excluded zero. Pre-infestation crown architecture did not credibly predict subsequent colonisation. These patterns are more consistent with persistent infestation imposing lasting architectural legacies than with strong pre-existing architectural susceptibility, suggesting rising liana abundance may progressively reduce forest stature and canopy density, with implications for tropical carbon storage projections.

## Introduction

Tropical forests represent the most carbon-dense and biodiverse terrestrial ecosystems on Earth, storing over half of all terrestrial carbon and harbouring at least two-thirds of known plant species (Malhi et al., 2014; Pan et al., 2011). Their structural complexity underpins these functions by governing light interception, competitive hierarchies, and aboveground biomass accumulation. Anthropogenic pressures, including land-use change, fragmentation, and climate-driven shifts in drought and disturbance regimes, are reshaping canopy organisation and species composition across the tropics (Laurance et al., 2014). One observed shift is the widespread increase in the abundance and biomass of lianas (Rueda-Trujillo et al., 2024; Schnitzer and Bongers, 2011). On Barro Colorado Island (BCI), liana stem density has increased by nearly one-third over a single decade (Schnitzer et al., 2021), with longer-term surveys documenting a 75% increase over 30 years alongside an 11.5% decline in tree density (Schnitzer et al., 2012). As liana abundance increases across the tropics (Rueda-Trujillo et al., 2024), resolving its impacts on host trees becomes essential for predicting future changes in forest structure and ecosystem function.

Lianas are woody vines that act as structural parasites, using host trees for mechanical support to access the canopy without investing in self-supporting tissue (Stevens, 1987; Stewart and Schnitzer, 2017). Lianas have high leaf-to-wood ratios, enabling rapid vertical and lateral expansion (Schnitzer et al., 2005). Lianas compete directly with hosts for light, water, and nutrients, and liana-infested trees have lower growth, reproduction, and survival than uninfested trees (Dias et al., 2017; Ingwell et al., 2010; Schnitzer et al., 2005). Infested trees are also shorter and less slender than uninfested trees (Schnitzer et al., 2005; Dias et al., 2017; Krishna Moorthy et al., 2026). Liana-removal experiments provide further evidence that reduced tree stature reflects a causal effect of lianas, with liana-infested trees remaining shorter for a given diameter than trees in liana-removal plots (Cao et al., 2025). At the forest level, the Gigante Peninsula liana-removal experiment, adjacent to BCI, found that lianas can reduce annual carbon uptake by up to 76% and shift production from woody biomass towards short-lived foliage (van der Heijden et al., 2015). Together, these findings demonstrate that lianas can alter individual-tree architecture and reduce forest carbon accumulation and biomass storage, highlighting potential links between liana-driven changes in tree structure and broader consequences for forest carbon dynamics.

Despite the broader implications of increasing liana abundance, whether liana infestation is primarily a cause or consequence of variation in tree architecture remains unresolved at the individual-tree level. Lianas can impose mechanical loads and competitive suppression on their tree hosts, hypothesised to result in structural changes consistent with chronic redirection of carbon allocation (Cao et al., 2025; Dias et al., 2017; Schnitzer et al., 2005). In the other direction, the probability of liana infestation is shaped by species-level traits and demographic processes. Variation in infestation prevalence across species is strongly associated with differences in liana shedding rates and infestation-induced mortality, both linked to shade tolerance: light-demanding species shed lianas more rapidly and suffer higher mortality when infested, resulting in lower prevalence, whereas shade-tolerant species accumulate lianas over time (Visser et al., 2018b, 2018a). However, colonisation rates vary little among tree species and are not clearly related to shade tolerance, indicating that additional factors determine which individual trees become infested. Crown architecture has long been hypothesised as one such factor: rapid monopodial growth and branch shedding may limit attachment, whereas shorter trees with wider, laterally connected crowns increase opportunities for vertical access and crown-to-crown transfer (Putz, 1984; Schnitzer et al., 2005; Visser et al., 2018a). Most studies, however, have lacked the combination of long-term infestation histories and high-resolution structural measurements required to disentangle whether architectural differences precede infestation or emerge as a consequence of persistent liana load.

Terrestrial laser scanning (TLS) now improves the structural resolution at which we can investigate structural feedbacks between trees and their liana hosts. TLS generates high-resolution three-dimensional point clouds, enabling extraction of metrics that link tree structure directly to ecological processes (Moorthy et al., 2018; Calders et al., 2020; Maeda et al., 2025). BCI offers a uniquely powerful context for applying these methods: the 50-ha Forest Dynamics Plot provides individual-level species identities, size measurements and spatial coordinates, while annual dendrometer surveys have recorded liana infestation scores for over a decade (Schnitzer et al., 2021). Combining a 2019 TLS structural snapshot with liana infestation histories spanning 2011–2025 provides a pre-scan infestation history, a point-in-time measure of tree structure, and post-scan infestation trajectories, enabling the directional pathways of the structural feedback to be separated. Few prior studies have combined longitudinal infestation records, individual-level three-dimensional structure, and a sample spanning the multi-year dynamics required to resolve this directionality. Cross-sectional studies can show that infested trees are shorter but cannot determine whether lianas suppressed host growth or whether shorter trees were simply more susceptible to colonisation (Moorthy et al., 2018, 2022).

To disentangle whether liana infestation primarily reflects a cause or a consequence of variation in tree architecture, we test two hypotheses. First, (H1, Figure 1) a history of persistent liana infestation produces measurable structural legacies in host tree architecture, such that persistently infested individuals exhibit lower height, crown area, crown volume and crown depth relative to trees that have never been infested or have lost their lianas. Lower height is interpreted as reduced or stagnant vertical growth rather than a decline from the tree’s original height, reflecting sustained mechanical loading and competitive light interception that redirect carbon allocation towards trunk diameter growth and constrain vertical and lateral crown development (Cao et al., 2025; Dias et al., 2017; Schnitzer et al., 2005). Second, (H2) intrinsic variation in tree architecture predicts subsequent liana infestation risk, with shorter trees and larger or deeper crowns expected to be more likely to become infested, because such architectures may facilitate liana establishment and accessibility through vertical access and lateral transfer between neighbouring crowns (Putz, 1984; Schnitzer et al., 2005; Visser et al., 2018a). By separating these directional processes, this framework allows the two directions of association between infestation history and tree structure to be evaluated separately.

**Figure 1.**
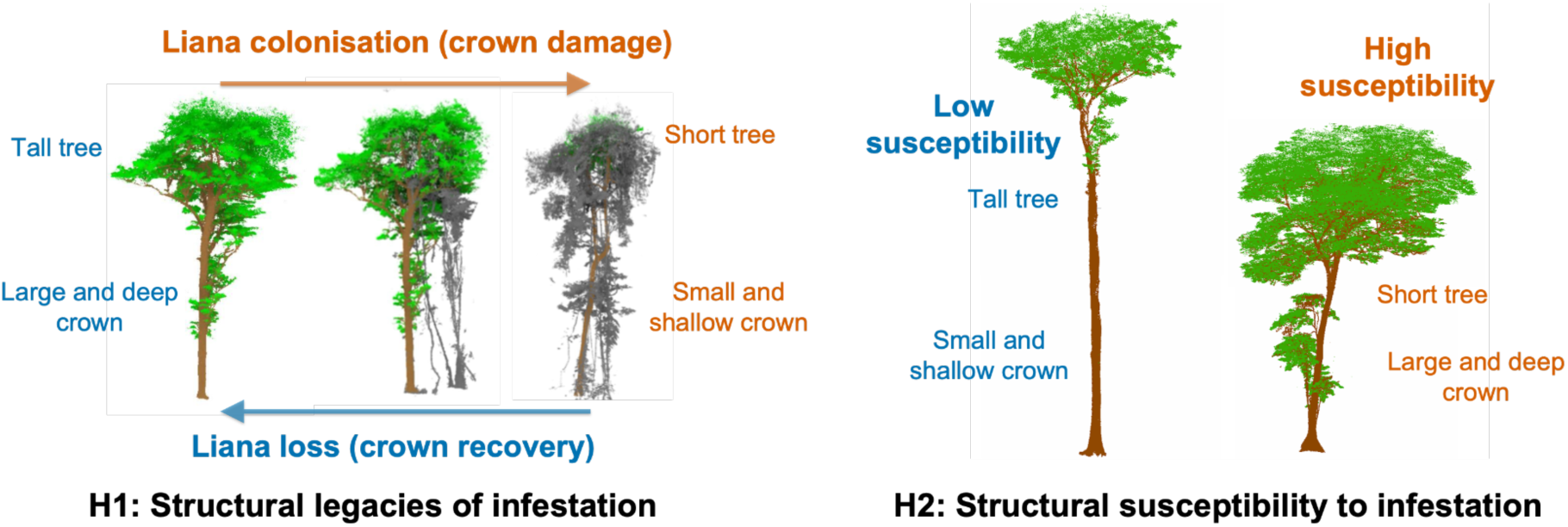
Conceptual framework for testing reciprocal structural feedbacks between liana infestation and host tree architecture. The framework tests whether persistent liana infestation is associated with structural legacies in host trees (H1), including reduced height and crown dimensions. Alternatively, it tests whether pre-existing tree architecture predicts subsequent infestation susceptibility (H2), with shorter trees and larger, deeper or more connected crowns expected to have greater infestation risk.

## Methods

### Study site

Barro Colorado Island (BCI), Panama (9°09′N, 79°51′W), is a 15.6 km² island in Gatun Lake supporting predominantly old-growth seasonally moist lowland tropical forest (Figure 2), with a pronounced dry season and mean annual rainfall of ∼2,600 mm. A 50-ha ForestGEO Forest Dynamics Plot was established on the central plateau in 1981; all trees ≥1 cm diameter were mapped, tagged, measured and identified, and the plot has been re-censused at approximately five-year intervals since 1982 (Condit, 1998). Lianas are surveyed separately in dedicated liana censuses (Schnitzer et al., 2012, 2021), making it a key system for studying liana–tree interactions.

**Figure 2.**
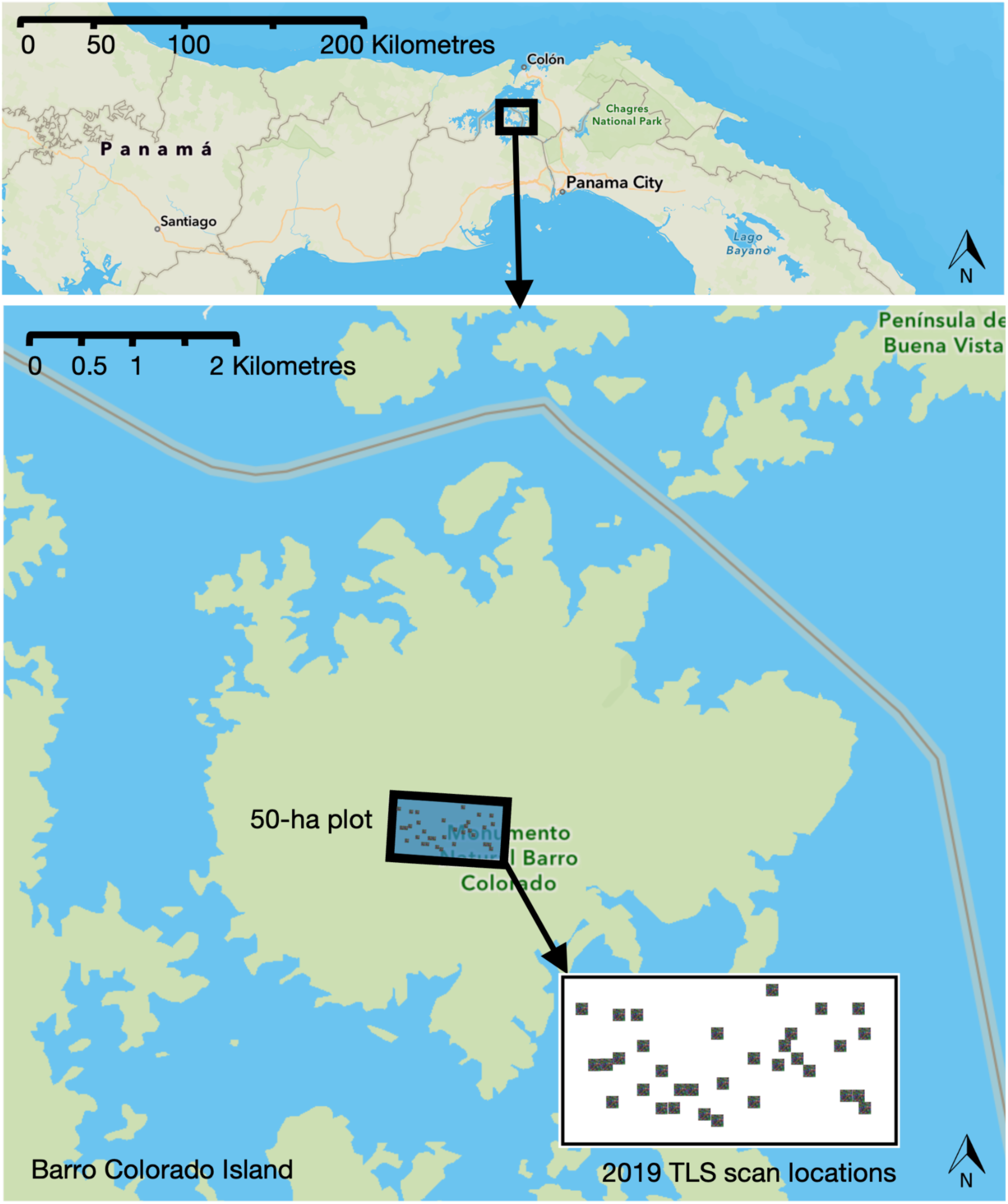
Location of the Barro Colorado Island (BCI) 50-ha Forest Dynamics Plot, Panama, and the subplots used for 2019 terrestrial laser scanning (TLS). The upper map shows the position of BCI within Panama, while the lower map shows the 50-ha plot within Gatun Lake and the distribution of TLS scan subplots. The inset shows the spatial arrangement of 2019 TLS scan locations. Base map and plot boundary data were adapted from the BCI 50-ha feature layer published by the GIS STRI Portal, Smithsonian Tropical Research Institute, licensed under a CC BY-SA 4.0.

### Long-term liana records

The liana records used in this study derive from 100 non-overlapping, spatially and size-stratified dendrometer subplots (Ramos et al., 2022). The dendrometer subplots are 40 × 40 m, nested within the 50-ha plot. Dendrometer census 8 (2011) was the first to record liana load in the crown of each surveyed tree, on a five-point scale (0 = no lianas, 1 = 1-25%, 2 = 26-50%, 3 = 51-75%, 4 = 76-100% of crown infested). Annual observations are made from the ground, using binoculars. In subsequent analyses this was transformed into three classes (0 = no lianas, 1 = 1-50%, light load, 2 = 51-100%, heavy load).

To reconstruct the individual-level infestation histories required to test H1 and H2, we merged annual dendrometer records across census years into a continuous liana infestation trajectory for each tree spanning 2011–2025, and subsetted these to the relevant window for each hypothesis (2011–2019 for H1; 2019–2025 for H2). Trees were classified as infested in any year with a non-zero liana score; when two censuses occurred within a year (2011–2018), the first was retained, which is unlikely to bias presence classifications because liana cover changes little within a season. For each individual we derived the first and most recent years of observed infestation and the total number of observed infested years, and assigned trees to five mutually exclusive infestation-history categories: never infested (no infestation 2011–2019; n = 62), recently infested (first infested 2016–2019 and infested in 2019; n = 12), persistently infested (first infested ≤ 2015 and infested in 2019; n = 99), recently lost lianas (last infested 2016–2018; n = 44) and lost lianas earlier (last infested ≤ 2015; n = 46). Individuals with insufficient pre-2019 records to classify infestation history were excluded. To increase statistical power, the two lost-lianas groups were pooled and the recently infested trees group (the smallest group) was removed, yielding three categories for H1 analyses. For H2 analyses, we restricted the sample to trees that were uninfested in 2019 and classified individuals into two mutually exclusive subsequent-colonisation categories: remained uninfested (no infestation recorded from 2019–2025; n = 112) and gained infestation (first observed with lianas after the 2019 census; n = 69). This classification isolates subsequent colonisation from prior infestation history, allowing pre-infestation architecture to be compared between trees that did and did not become infested.

### TLS data acquisition and processing

To maximise crown coverage and minimise occlusion artefacts, TLS data were collected in 24 fixed 40 × 40 m subplots using a RIEGL VZ-400 terrestrial laser scanner. Each subplot was scanned from 25 positions distributed across a 60 × 60 m area centred on the subplot, with scan positions spaced approximately 15 m apart. Further details of the field sampling design and TLS acquisition protocol are provided in Krishna Moorthy et al. (2026). TLS data were co-registered and merged from all scan positions into a single three-dimensional point cloud for each of the 24 subplots using RiSCAN Pro (v2.5.3; Wilkes et al., 2017). From these registered point clouds, trees ≥ 20 cm diameter at breast height (DBH) were segmented. Analyses were restricted to trees at or above this size because lianas preferentially infest larger, canopy-accessing individuals, where increased light availability facilitates establishment and persistence (Schnitzer et al., 2005).

We processed each co-registered subplot point cloud through a three-stage segmentation pipeline to isolate individual tree crowns for architectural analysis (Figure 3). Candidate trees were first extracted from the full subplot point clouds using RayCloudTools (Lowe and Stepanas, 2021), which performs robustly in structurally complex forest environments (Cherlet et al., 2026). A height filter retaining only objects taller than 5 m was applied to remove understorey vegetation. TLS point clouds were subsequently aligned to the ForestGEO stem map, and segmented trees were matched to tagged field individuals using spatial position and DBH, with known species identity and tree morphology used as additional visual cues where needed. Verified trees were then cleaned in CloudCompare to remove residual liana material and segmentation artefacts prior to architectural analysis.

**Figure 3.**
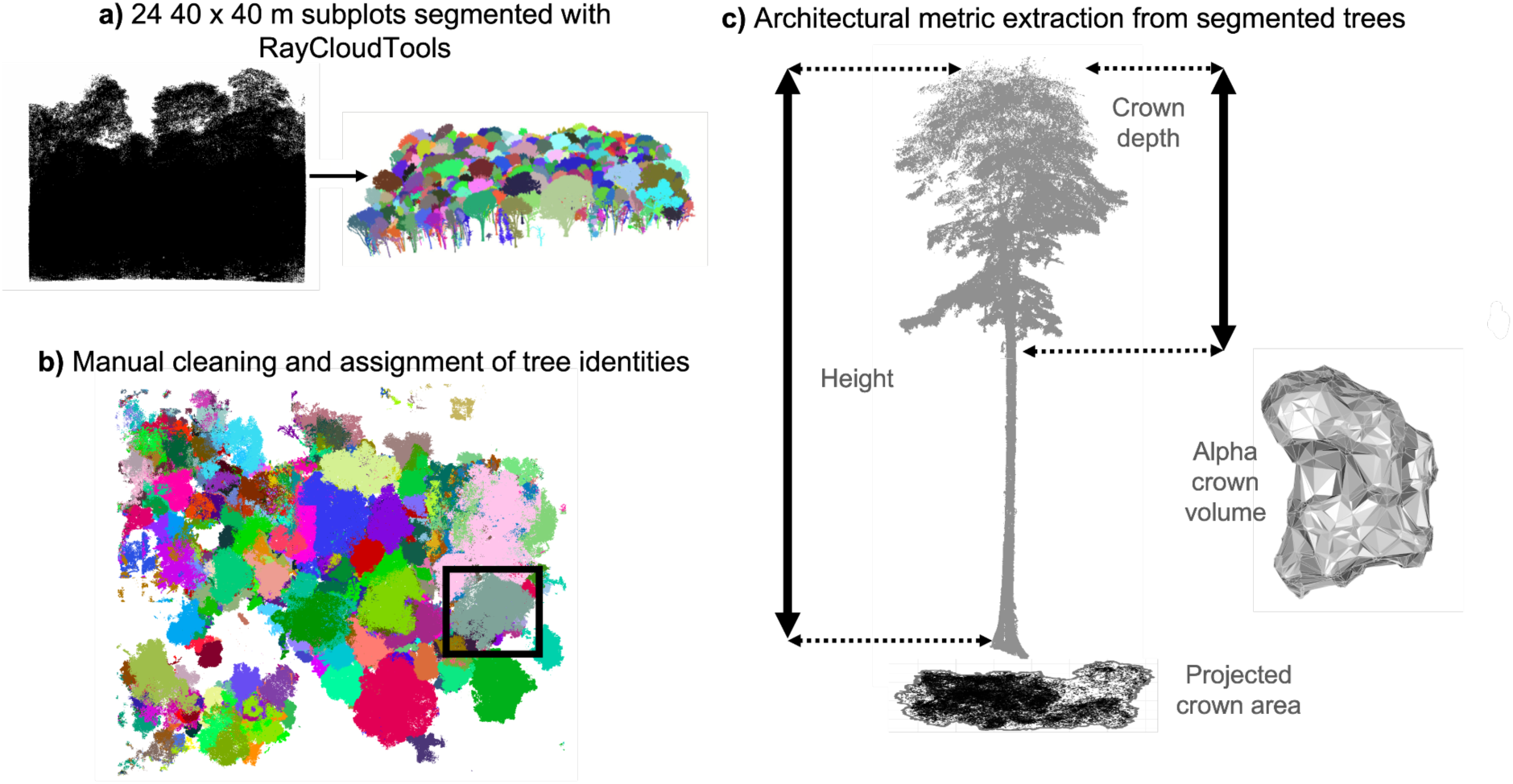
Workflow used to transform raw TLS point clouds from the Barro Colorado Island 50-ha plot into cleaned individual trees for structural analyses. (a) 24 co-registered subplot scans segmented using RayCloudTools (Lowe and Stepanas, 2021). (b) Manual cleaning of outputs, coordinate transformation to the stem map, and assignment to tagged individuals with associated liana records. (c) Extraction of tree height, projected crown area, crown volume and crown depth from cleaned point clouds using ITSMe R package (Terryn et al., 2023).

### Architectural metrics

Tree height, projected crown area, crown volume, and crown depth were calculated using the ITSMe R package (v1.0.0; Terryn et al., 2023) via the basicsummary() function. The crown was defined as all points located above the first major branch, thereby excluding the main stem and lower bole, and buttress detection was applied to prevent buttress points from being incorrectly classified as crown material. The four metrics capture distinct dimensions of tree architecture with direct ecological relevance. Projected crown area was estimated as the area enclosed by a two-dimensional concave hull fitted to the crown point cloud (alpha = 2), quantifying horizontal canopy expansion, serving as a proxy for competitive reach and light interception footprint. Crown volume, the three-dimensional alpha-shape volume of the same point cloud (alpha = 1), integrates both horizontal and vertical extents to capture the total canopy space occupied by the tree (Terryn et al., 2023). Crown depth, the vertical distance between the highest and lowest crown points, represents the vertical extent of the crown and its potential for vertical light capture. Tree height captures total stature, with direct implications for canopy position and competition for light (Cao et al., 2025; Martínez Cano et al., 2019).

### Field data collection

To extend H2 analyses of pre-infestation tree stature beyond the TLS dataset, we used ground-measured tree heights collected in 2011 with laser rangefinders (Larjavaara and Muller-Landau, 2013). To characterise liana colonisation pathways and climbing modes, we revisited the subplots in 2025 and recorded climbing mode(s) of associated lianas for trees of the eight most abundant species (≥10 individuals per species), following Sperotto et al. (2020), together with infestation status (none, direct, lateral, both). These descriptive observations provide context for the colonisation analyses but are not central to the structural tests.

## Statistical analysis

We conducted all statistical analyses in R (version 4.5.2; R Core Team, 2025), using the brms package (Bürkner, 2017). TLS-derived crown structural metrics (2019) were joined with individual-level metadata from three sources: long-term dendrometer records of liana infestation score and DBH (2011–2025), estimates of percentage liana crown cover (2019), and field observations of climbing mode and infestation status (2025). DBH and all structural metrics were log-transformed to linearise allometric relationships (Cao et al., 2025; Martínez Cano et al., 2019). Species and 40 × 40 m subplot were included as random intercepts in the models to account for taxonomic variation in crown form and spatial autocorrelation in habitat and liana distributions, respectively (Bai et al., 2022; Moorthy et al., 2022).

To test whether persistent liana infestation leaves measurable structural legacies (H1), Bayesian multilevel models were fitted for each structural metric, with the liana infestation history category and log(DBH) as fixed effects and species and subplot as random intercepts. We fitted Bayesian multilevel models in the form of Equation 1:

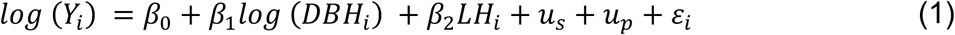

where *Y_i_* is the structural metric for individual *i*, *LH* is liana infestation history (never infested, lost lianas, persistently infested), *DBH_i_* is trunk diameter at breast height (cm), *β*0 , *β*1, and *β*2 are fixed effect coefficients, *u_s_* and *u_p_* are species- and subplot-level random intercepts, and *ε_i_* is the residual error. Effects are reported as posterior means with 95% credible intervals (CrI), back-transformed to approximate percentage differences where appropriate; effects were considered credible where the 95% CrI excluded zero.

To test whether pre-infestation crown architecture predicts subsequent colonisation (H2), a separate Bayesian multilevel model was fitted for each architectural metric. Rather than modelling infestation outcome as the response, each pre-infestation structural metric was modelled as the response, with subsequent infestation outcome included as a fixed effect. This tests whether trees that subsequently gained infestation differed in pre-existing structure from trees that remained uninfested, after accounting for DBH:

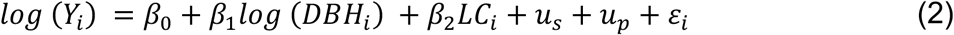

where *LC_i_* denotes subsequent liana colonisation outcome, with two mutually exclusive categories: gained infestation (n = 69) and remained uninfested through 2025 (n = 112); all other terms are defined above.

To further test H2 using a larger dataset of pre-infestation tree stature, we analysed 2011 ground-measured tree heights (n = 414) using asymptotic height– diameter models. We compared log–log and Michaelis–Menten formulations using WAIC and used the best-supported formulation for inference. The selected formulation was consistent with previous work showing that saturating generalised Michaelis– Menten models provide the best fit to tree height–diameter relationships at BCI (Martínez Cano et al., 2019). The model allowed the asymptotic height, scaling exponent and half-saturation diameter to vary by liana colonisation outcome:

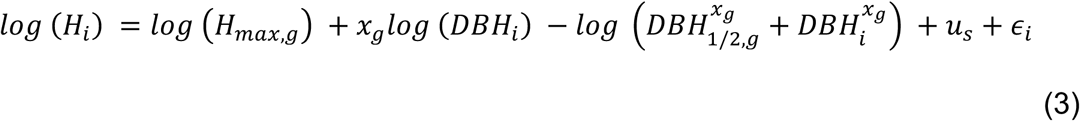

where *H_i_* is tree height (m) for individual *i*, *H_max_*_,g_is the asymptotic tree height for liana colonisation outcome group *g*, *DBH*_1/2,g_ is the diameter at breast height at which predicted height reaches half the asymptotic value, and *x*_g_ is the scaling exponent. Full model specifications are provided in Supporting Information Table S1.

A slenderness index (height/DBH) and crown base height ratio (crown base height / total tree height) were also evaluated as composite predictors.

## Results

### Structural legacies of infestation (H1)

We found partial support for H1 (Figure 4), which predicted that persistent liana infestation would leave measurable architectural legacies: persistently infested trees showed credibly reduced height, crown projected area, and crown volume relative to never-infested individuals, though crown depth and slenderness did not differ credibly. Tree height showed credible negative posterior estimates for both infestation categories relative to never-infested individuals. Persistently infested trees were ∼11.0% shorter (95% CrI: −14.7 to −7.2%, see Figure 4), while trees that had lost lianas were ∼5.0% shorter (95% CrI: −8.6 to −1.2%). Trees in the lost-lianas category had substantially lower peak infestation loads (median: score 1, 1–25% crown cover) and shorter infestation durations (median: two years) than persistently infested individuals (median: score 4, 76–100%; median: eight years; Figure S2). Persistently infested trees also encompassed a broader range of infestation intensities, whereas the lost-lianas category was disproportionately represented by lower infestation loads. Crown projected area showed a credible reduction only in the persistently infested category (−22.3%; 95% CrI: −33.4 to −10.0%), while the lost-lianas group did not credibly differ from never-infested trees (−10.1%; 95% CrI: −22.1 to 3.5%). Crown volume followed the same pattern: a credible reduction for persistently infested individuals (−25.9%; 95% CrI: −40.2 to −9.1%), and no credible effect for trees that had lost lianas (−7.4%; 95% CrI: −24.4 to 12.8%). Slenderness did not differ credibly between infestation categories and the never-infested reference once species identity was accounted for (persistently infested: −7.0%, 95% CrI: −14.9 to 0.4%; lost lianas: −3.4%, 95% CrI: −10.9 to 3.9%). Crown depth showed no credible difference for either category (persistently infested: −5.5%, 95% CrI: −13.7 to 3.6%; lost lianas: ∼0%, 95% CrI: −8.6 to 9.4%), nor did crown base height ratio (persistently infested: −7.1%, 95% CrI: −17.4 to 4.3%; lost lianas: −4.0%, 95% CrI: −13.8 to 6.9%). Combined, these results indicate that persistent infestation is associated with reduced realised height and crown size. Although crown depth and crown base height ratio showed no credible differences, point estimates were negative and effects in this direction cannot be ruled out.

**Figure 4.**
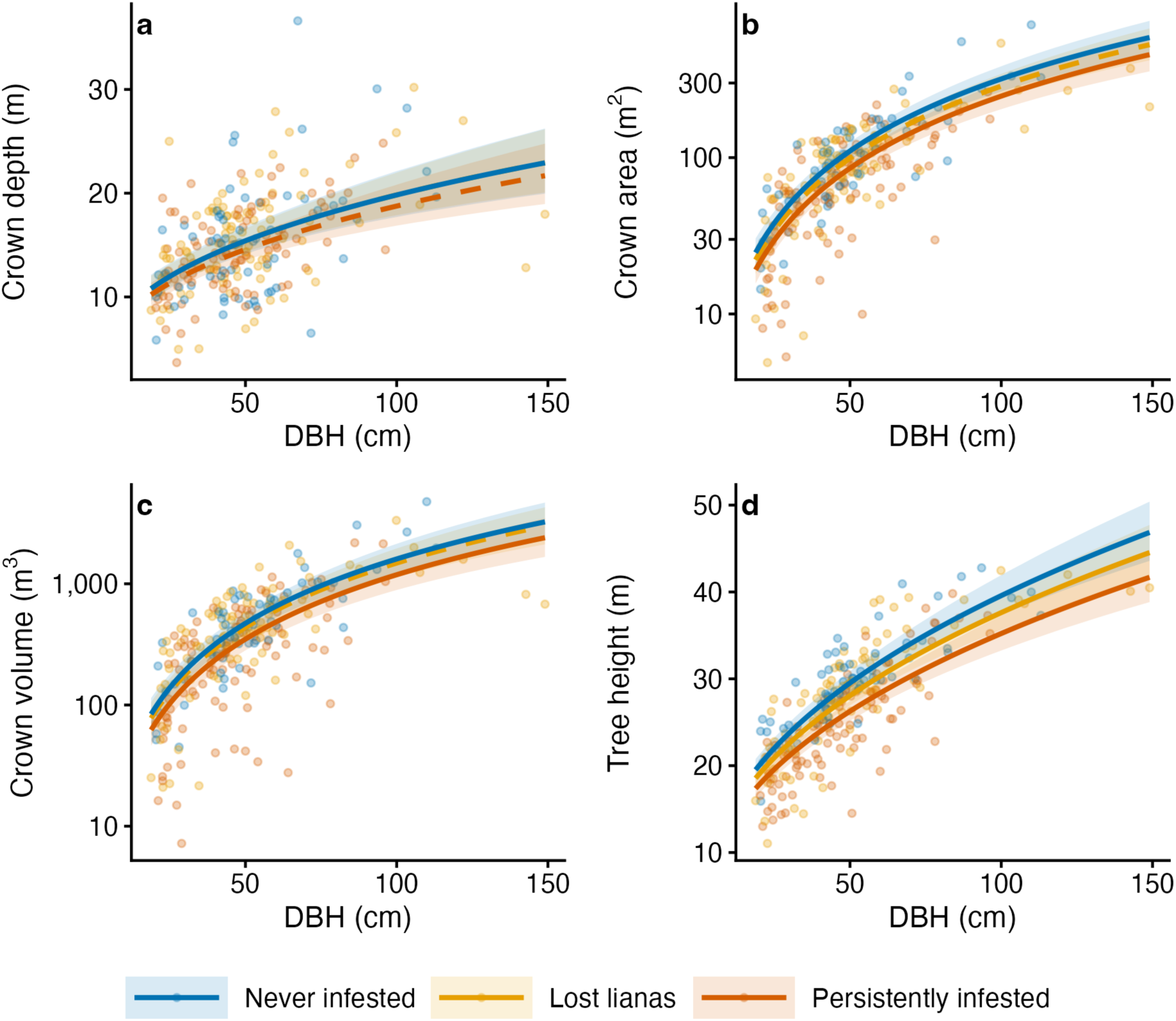
Persistent liana infestation is associated with credibly reduced tree height, crown projected area, and crown volume relative to never-infested trees, while trees that lost lianas between 2011 and 2019 occupy an intermediate structural position. Scatter plots show allometric relationships between trunk diameter at breast height (DBH, cm) and tree architectural metrics on their original scales; models were fitted using log-transformed DBH, and (a) crown depth (m), (b) crown projected area (m²), (c) crown volume (m³), and (d) tree height (m) for 251 trees across 67 species, classified into three mutually exclusive infestation-history categories based on liana load data from 2011: never infested (blue), trees that lost lianas before the 2019 TLS scan (orange), and persistently infested trees (first infested before 2015 and still infested in 2019; red). Points represent individual trees and lines show Bayesian posterior population-level fits from brms Student-t multilevel models that include log(DBH) as a fixed-effect covariate and random intercepts for species and subplot; ribbons show 95% credible intervals. Solid lines indicate effects whose 95% credible interval excludes zero, and dashed lines indicate non-credible effects.

**Figure 5.**
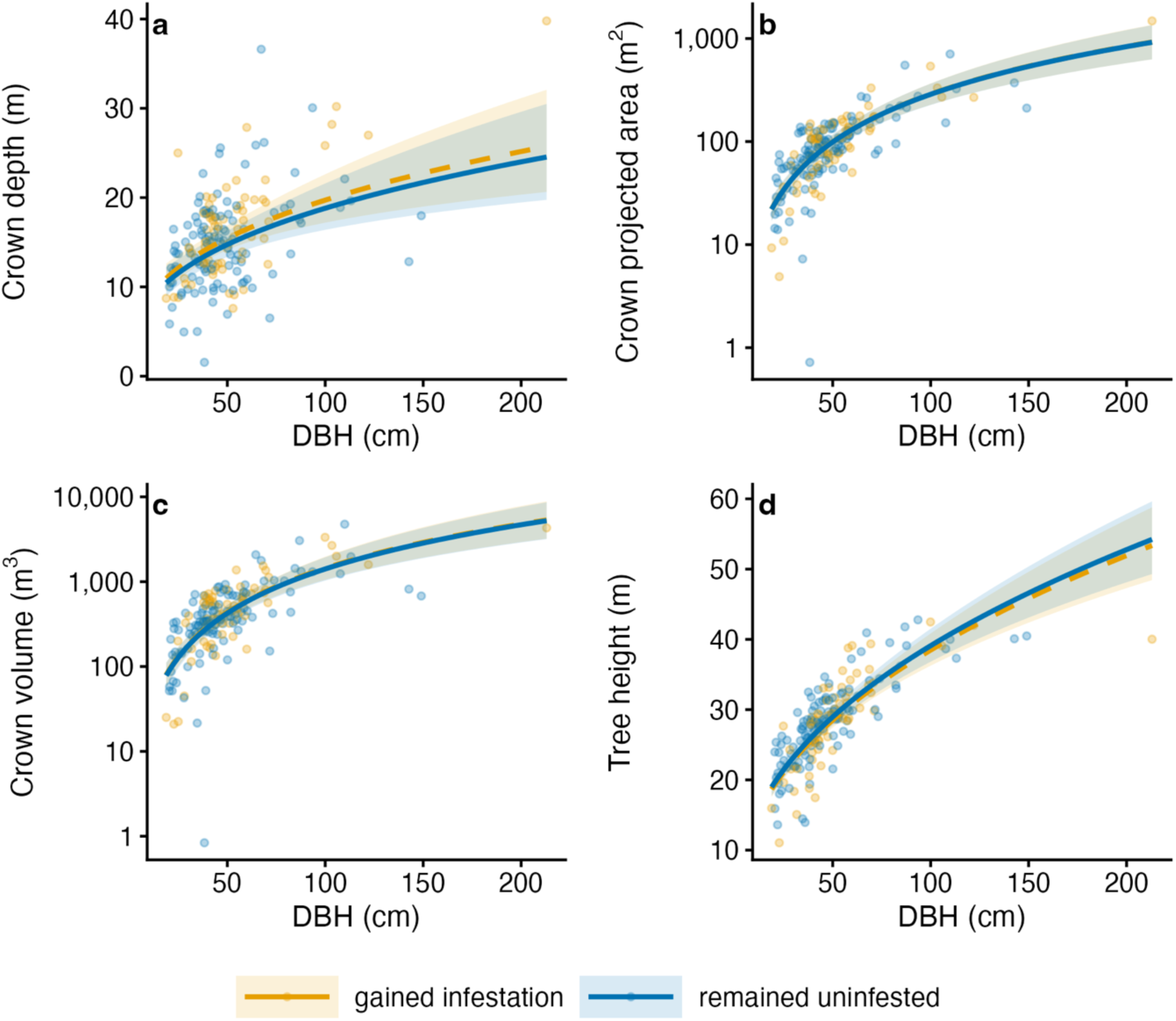
No TLS-derived structural metric showed a credible difference between trees that did and did not subsequently gain lianas (2019-2025), with all 95% credible intervals spanning zero. Scatter plots show log-log allometric relationships between trunk diameter at breast height (DBH, cm) and (a) crown depth (m), (b) crown projected area (m²), (c) crown volume (m³), and (d) tree height (m) for 181 trees across 61 species that were uninfested in 2019, classified by subsequent infestation outcome: trees that remained uninfested by 2025 (blue) and trees that gained liana infestation by 2025 (orange). Points represent individual trees and lines show Bayesian posterior population-level fits from brms Student-t multilevel models that include log-transformed DBH as a fixed-effect covariate and random intercepts for species and subplot; ribbons show 95% credible intervals.

### Structural determinants of infestation (H2)

In contrast, we found no support for H2, which predicted that shorter trees with larger or more structurally accessible crowns would show greater susceptibility to liana colonisation. Posterior estimates for the effect of gaining infestation relative to remaining uninfested were small and centred near zero across all metrics (height: −1.0%, 95% CrI: −4.6 to 2.5%; crown projected area: +2.3%, 95% CrI: −8.9 to 14.8%; crown volume: +3.0%, 95% CrI: −12.1 to 20.8%; crown depth: +1.6%, 95% CrI: −6.0 to 9.9%), with all 95% credible intervals spanning zero. Trees that subsequently gained infestation did not show credibly lower pre-existing slenderness than those that remained uninfested (−4.1%, 95% CrI: −11.4 to 2.8%).

When H2 was extended to the larger 2011 height dataset, predicted heights were slightly lower for trees that subsequently gained infestation across the size range examined (25–150 cm DBH: −2.4 to −8.0%), but credible intervals overlapped at all sizes, providing no conclusive evidence that shorter trees are more susceptible to infestation (Fig. S1).

## Discussion

Increasing liana abundance represents one of the most widespread compositional shifts occurring across tropical forests (Rueda-Trujillo et al., 2024; Schnitzer and Bongers, 2011), but the long-term structural consequences of this shift for host trees remain poorly resolved. Although previous studies have shown that heavily infested trees are often shorter and possess smaller crowns, most have relied on cross-sectional comparisons or short-term experimental manipulations that cannot distinguish whether lianas actively suppress tree architecture or simply colonise trees with pre-existing structural characteristics that increase susceptibility (Cao et al., 2025; Loubota Panzou et al., 2022; Dias et al., 2017; Moorthy et al., 2022; Krishna Moorthy et al., 2026). Previous work has not directly examined whether persistent infestation leaves cumulative architectural legacies or whether host trees structurally recover following liana loss. Resolving these directional pathways is critical because chronic liana infestation has the potential to alter canopy structure, competitive dynamics, and forest carbon storage at ecosystem scales (Schnitzer and DeFilippis, 2025; van der Heijden et al., 2015).

By combining longitudinal individual-level infestation histories with high-resolution three-dimensional structural measurements, we disentangle the reciprocal relationship between liana infestation and tree architecture in a way that has not previously been possible. We show that persistent infestation was consistently associated with reduced tree stature and crown size, whereas crown architecture of liana-free trees showed little difference among trees that were and were not subsequently colonized. Together, these findings support the conclusion that persistent liana infestation contributes to long-term architectural shifts, and do not support the hypothesis that host crown architecture strongly influences infestation probability.

The strong reductions in host tree height and crown size associated with persistent infestation are consistent with a growing body of evidence that lianas alter tree architectural development (Cao et al., 2025; Dias et al., 2017; Krishna Moorthy et al., 2026; Meunier et al., 2021; Schnitzer et al., 2005; van der Heijden et al., 2015). Recent analyses from the same BCI 50-ha plot extend this pattern beyond tree height, showing that heavy liana loads are associated with substantially reduced crown area and crown volume (Krishna Moorthy et al., 2026). Experimental liana-removal studies on Gigante Peninsula demonstrated that lianas reduce forest-level carbon accumulation and woody productivity by constraining host growth (van der Heijden et al., 2015), while TLS-based analyses from the same experimental system showed that liana-infested trees are shorter for a given diameter and therefore likely store less carbon than uninfested individuals (Cao et al., 2025). The stronger reduction in tree stature observed here compared with the Gigante Peninsula experiment (Cao et al., 2025) may reflect the different scales at which liana effects were quantified. The Gigante Peninsula is also a secondary forest, which may differ from the old-growth forest studied here in the prevalence and persistence of liana infestation. Plot-level treatment contrasts include both heavily and lightly infested trees, whereas the present study isolates long-term structural differences associated with persistent infestation histories at the individual-tree level. More broadly, global analyses of tree crown architecture suggest that crown dimensions and height scaling relationships are shaped by environmental constraints and competitive pressures that govern carbon allocation between vertical growth and lateral canopy expansion (Jucker et al., 2025). Within this broader framework, persistent liana infestation may represent an additional biotic constraint capable of shifting host trees away from the crown scaling relationships expected under a given set of environmental conditions.

The intermediate structural profile of trees that had lost lianas could be interpreted as evidence for architectural recovery following liana loss, but may also be explained by lower cumulative infestation loads and shorter infestation durations. Indeed, their weaker structural shifts are more parsimoniously explained by reduced cumulative liana burden. This distinction matters because previous studies have examined architecture in relation to current liana load or long-term experimental treatment, but have not separated the effects of an individual tree’s infestation history from its current infestation state (Cao et al., 2025; Krishna Moorthy et al., 2026). Our findings instead suggest that long-term structural suppression depends not only on contemporary liana load, but also on the duration and cumulative history of infestation. The persistence of reduced crown dimensions in chronically infested trees may therefore have functional consequences beyond architecture alone, because degraded crown condition is strongly associated with reduced growth and elevated mortality risk in tropical forests (Arellano et al., 2019). Although some trees may partially recover following liana loss, our results suggest that persistent infestation can leave long-lasting architectural legacies, and it remains unclear whether heavily infested individuals ever fully recover their original crown structure. Variation among species in the ability to shed lianas or tolerate chronic crown occupation may therefore play an important role in determining long-term structural trajectories following infestation, with potential consequences for canopy function and forest carbon cycling.

In contrast to the strong architectural legacies associated with persistent infestation in H1, pre-infestation architecture showed little evidence of strongly determining subsequent colonisation risk under H2. This distinction is important because one alternative explanation for the reduced height observed in persistently infested trees is that lianas preferentially colonise trees that are already shorter and more structurally accessible, rather than actively suppressing growth following infestation. Previous work has long hypothesised that host tree structure influences liana susceptibility, with shorter trees, deeper crowns, and greater lateral canopy connectivity potentially facilitating vertical access and crown-to-crown transfer (Putz, 1984; Visser et al., 2018b). However, our results provide limited support for a strong architectural predisposition, although the absence of credible effects may reflect insufficient power to detect small but genuine architectural differences. This interpretation is consistent with work demonstrating that long-term differences in infestation prevalence among species on BCI are driven primarily by variation in liana shedding and infestation-induced mortality rather than colonisation itself (Visser et al., 2018b). Collectively, these findings suggest that the processes governing liana persistence following establishment may be more important than initial colonisation in shaping long-term infestation patterns.

The weak predictive power of crown architecture in H2 suggests that neighbourhood and spatial processes may dominate liana establishment dynamics at local scales. In addition to tree species, subplot was included as a random effect to account for habitat and broad-scale spatial clustering of liana sources. However, fine-scale neighbourhood proximity, such as adjacency to heavily infested trees or canopy gaps, could not be controlled for and likely explains additional residual variation in colonisation outcomes. Liana distributions are often strongly spatially aggregated, reflecting dispersal limitation, local source pools, and canopy disturbance dynamics rather than host-tree properties alone (Bai et al., 2022; Schnitzer et al., 2021). On BCI, local canopy disturbance promotes rapid liana proliferation through clonal expansion and increased canopy access within gaps (Schnitzer et al., 2021), while broader theoretical frameworks increasingly describe liana spread as an epidemiological process governed by interactions among infestation force, host demography, and forest structure (De Deurwaerder et al., 2024; Muller-Landau & Pacala, 2020). Within this context, the absence of strong crown-metric predictors in H2 suggests that whether a tree becomes infested may depend less on its intrinsic architecture than on its spatial exposure to nearby liana sources and disturbed canopy environments. This interpretation is also consistent with the 2025 field observations, which showed frequent direct ascent and substantial use of active climbing mechanisms, although many newly infested trees were colonised through both direct and lateral pathways (Figures S8-S10). Nevertheless, both the TLS and ground-based height analyses showed weak directional trends towards shorter stature in trees that subsequently gained infestation, indicating that small pre-existing structural differences cannot yet be ruled out entirely.

Species differences in tolerance of liana infestation and in liana shedding likely mediate how strongly chronic infestation translates into long-term architectural suppression. Species-level random intercepts (Figures S3–S7) revealed systematic variation in crown metrics not explained by infestation history or tree size, consistent with the established role of traits like bark texture, branch-shedding capacity, and shade tolerance in governing liana persistence and host vulnerability (Putz, 1984; Visser et al., 2018a, 2018b). This is consistent with Krishna Moorthy et al. (2026), who found substantial interspecific variation in baseline crown area and crown volume within the same BCI forest. The escape hypothesis predicts that tall, fast-growing trees with rapid stem thickening and branch shedding can avoid or shed lianas more effectively (Putz, 1984), whereas shade-tolerant species may accumulate lianas due to reduced shedding capacity (Visser et al., 2018b). Previous work on BCI has demonstrated that liana shedding rate and infestation-induced mortality, rather than colonisation rate, drive interspecific variation in long-term infestation prevalence; shade-tolerant species tend to accumulate higher liana loads because they shed lianas less rapidly and experience lower mortality when infested (Visser et al., 2018b). The species-level variation captured by random intercepts, together with the null H2 result, suggests that taxonomic identity may be a stronger driver of infestation outcomes than the individual crown architecture metrics considered here. Other species-specific architectural traits not captured by the metrics examined here may also contribute to variation in infestation outcomes. Future modelling efforts could benefit from incorporating species-level shade tolerance, bark traits, and potentially branching patterns and other species-specific architectural traits (De Deurwaerder et al., 2024; Visser et al., 2018a).

Several limitations constrain the extent to which the rate and trajectory of architectural change can be resolved from the present dataset. Because this study relies on a single TLS snapshot, it cannot estimate how rapidly individual trees change structurally following infestation, nor resolve the extent or timing of recovery after liana loss. Sample sizes for some infestation-history categories were limited, necessitating category pooling and reducing power to detect finer-scale differences among infestation trajectories. Because liana species identity was unavailable, we treated all infestations as ecologically equivalent. Differences among liana species in growth strategy, leaf distributions within hosts, and more may nevertheless influence their architectural effects and could contribute to unexplained variation among trees (Ichihashi & Tateno 2011). Repeat TLS surveys of the same trees at approximately five-year intervals would therefore provide a practical approach for quantifying structural change, tracking recovery trajectories, and determining whether crown dimensions remain persistently suppressed or recover following liana loss (Calders et al., 2020).

Rising liana abundance across the tropics makes understanding its cumulative structural and carbon consequences a priority for forest ecology and carbon accounting. By combining long-term infestation histories with individual-level three-dimensional crown structure, this study provides a framework for disentangling the causes and consequences of liana infestation while highlighting the potential for persistent infestation to reshape tropical forest architecture over time. As tropical forests face accelerating pressures from climate change and global disturbance, such structural change may have important implications for the long-term stability of one of Earth’s largest carbon sinks.

## Supporting information

SOM

## Acknowledgements

We are grateful to the Smithsonian Tropical Research Institute for logistical support on Barro Colorado Island. We also thank the Crankstart Scholarship Programme and the Department of Biology, University of Oxford for financial support that enabled fieldwork in Panama in 2025. P. Ramos and P. Villareal collected the time series of liana loads and the tree height data on the dendrometer trees. J.M. and J.D. supported field work in 2025. Financial support for the dendrometer data collection was provided by Smithsonian ForestGEO and the HSBC Climate Partnership. TLS data collection was funded by the ForestGEO Research Grants Program of 2018. F.M. was supported by the Research Foundation – Flanders (FWO) as a senior postdoctoral fellow (Grant No 1214723N) and under an ERC runner-up project (G0BHJ26N). H.V. was supported by Research Foundation – Flanders (FWO) with grant number G002321N. RSG was supported by a NERC Pushing the Frontiers grant (NE/X013766/1).Co–senior authors S.M.K.M. and R.S.G. contributed equally to this article, and they can list themselves last in author order on their CVs and applications.

## Supporting Information

Additional Supporting Information may be found in the online version of this article. Figures S1-S10 and Table S1 are cited in the text.

## Author Contributions

R.Y., S.M.K.M., and R.S.G. designed the research. R.Y. performed the analyses and wrote the manuscript. H.C.M. contributed to conceptual design, data provision and analytical interpretation. F.M. and H.V. contributed to the TLS data collection and analysis. S.M.K.M. contributed to TLS data collection and processing, analytical interpretation, guidance and funding acquisition. R.S.G. contributed to conceptual guidance and manuscript revision. All authors contributed to writing the manuscript.

## Data Accessibility Statement

The R code used for statistical analyses, TLS processing and figure generation is available at: https://github.com/biologyanonymous/liana-project.git. TLS-derived individual-tree and subplot point clouds are large files and will be made available from the corresponding author on request, or deposited subject to repository constraints.

