## Supplementary material for "Long-term liana infestation leaves architectural legacies in host trees": SOM

**Article type:** Letter

**Running title:** Liana legacies in tree architecture

Rosie Young[1], Helene C. Muller-Landau[2], Félicien Meunier[3,4] , Hans Verbeeck[3], Sruthi M. Krishna Moorthy[1, 5, 6] * and Roberto Salguero-Gómez[1,6] *

[1] Department of Biology, University of Oxford, Oxford, UK

[2] Quantitative Forest Ecology Lab, Smithsonian Tropical Research Institute, Panama City, Panama

[3] Q-ForestLab, Department of Environment, Ghent University, Belgium

[4] Department of water and climate, Vrije Universiteit Brussel, Brussels, Belgium

[5] Environmental Change Institute, School of Geography and Environment, Oxford, UK

[6] Pembroke College, University of Oxford, Oxford, UK

* Shared senior authors

Supporting Information **1**

Model specifications and selection 2

Complementary height analysis 4

Peak infestation intensity vs. infestation duration 5

Random effects (species specific effects) 6

Climbing modes/ liana infestation pathway 11

**Model specifications and selection**

Structural metrics (tree height, crown projected area, crown volume, crown depth, and crown base height ratio) were fitted as Bayesian multilevel linear models using the brms package, implementing Hamiltonian Monte Carlo sampling (No-U-Turn Sampler) in Stan. All response variables and DBH were log-transformed prior to analysis. Fixed effects included liana infestation category and log-transformed DBH; varying intercepts for species and subplot accounted for interspecific variation and spatial clustering. Slenderness (height/DBH) was modelled using the same Bayesian multilevel model structure as the other metrics, but without log(DBH) as a fixed effect, since DBH is algebraically embedded in the ratio, making an additional size covariate redundant. Species and subplot were retained as random intercepts for consistency with all other models.

Model convergence was assessed using the potential scale reduction statistic ($\hat{R}$), effective sample sizes, and inspection for divergent transitions. All brms models achieved satisfactory convergence ($\hat{R} < 1.01$). Parameter estimates are reported as posterior means with 95% credible intervals, with effects interpreted as credibly different from zero where intervals do not overlap zero. Coefficients from log-transformed models were back-transformed to approximate percentage differences where appropriate.

**Table S1. Specifications of statistical models fitted to test hypotheses H1 and H2.** For H1, trees were classified from 2011–2019 liana records as never infested (no observed infestation during 2011–2019), lost lianas (previously infested but uninfested by the 2019 TLS survey), or persistently infested (first infested ≤2015 and still infested in 2019); recently infested trees were excluded from H1 analyses. For H2 TLS models, trees uninfested in 2019 were classified according to subsequent colonisation outcome as gained infestation by 2025 or remained uninfested through 2025. The 2011 height analysis used the corresponding subsequent-colonisation contrast. Bayesian multilevel linear models (LMMs) were fitted using the brms package with Hamiltonian Monte Carlo sampling in Stan. Species and subplot were included as random intercepts in TLS based analyses to account for interspecific variation in crown architecture and spatial clustering of liana infestation.

| H | Response variable | Model type | Fixed effects | Random effects | n obs | n species |
|---|---|---|---|---|---|---|
| **1** | log(Tree height) | brm LMM (Student-t) | log(DBH) + H1 Category | (1\|species) + (1\|subplot) | 251 | 67 |
| **1** | log(Crown projected area) | brm LMM (Student-t) | log(DBH) + H1 Category | (1\|species) + (1\|subplot) | 251 | 67 |
| **1** | log(Crown volume) | brm LMM (Student-t) | log(DBH) + H1 Category | (1\|species) + (1\|subplot) | 251 | 67 |
| **1** | log(Crown depth) | brm LMM (Student-t) | log(DBH) + H1 Category | (1\|species) + (1\|subplot) | 251 | 67 |
| **1** | Slenderness (height/DBH) | brm LMM (Student-t) | H1 Category | (1\|species) + (1\|subplot) | 251 | 67 |
| **1** | log(Crown base height ratio) | brm LMM (Student-t) | H1 Category | (1\|species) + (1\|subplot) | 251 | 67 |
| **2** | log(Tree height) | brm LMM (Student-t) | log(DBH) + H2 Category | (1\|species) + (1\|subplot) | 181 | 61 |
| **2** | log(Crown projected area) | brm LMM (Student-t) | log(DBH) + H2 Category | (1\|species) + (1\|subplot) | 181 | 61 |
| **2** | log(Crown volume) | brm LMM (Student-t) | log(DBH) + H2 Category | (1\|species) + (1\|subplot) | 181 | 61 |

| | | | | | | |
|---|---|---|---|---|---|---|
| **2** | log(Crown depth) | brm LMM (Student-t) | log(DBH) + H2 Category | (1\|species) + (1\|subplot) | 181 | 61 |
| **2** | Slenderness (height/DBH) | brm LMM (Student-t) | H2 Category | (1\|species) + (1\|subplot) | 181 | 61 |
| **2** | log(Crown base height ratio) | brm LMM (Student-t) | H2 Category | (1\|species) + (1\|subplot) | 181 | 61 |
| **2** | Height (2011) | brm NLM (Gaussian) | $H_{max,g}$ + $x_g$ + $DBH_{1/2,g}$ ~ H2 2011 Category | (1\|species) | 414 | 85 |

### Complementary height analysis

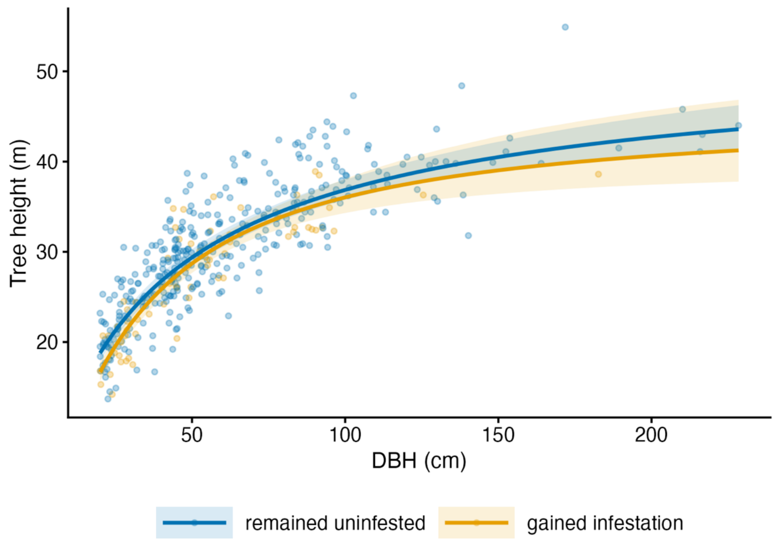


**Figure S1. Trees that subsequently gained liana infestation between 2011 and 2019 show no credible difference in asymptotic height relative to trees that remained uninfested, indicating that pre-existing tree structure does not robustly predict colonisation susceptibility.** The figure shows ground-measured tree height (m; 2011) as a function of trunk diameter at breast height (DBH, cm) for 414 trees across 85 species that were uninfested in 2011, classified by subsequent infestation outcome: trees that remained uninfested (blue) and trees that gained liana infestation by 2019 (yellow). Lines show posterior population-level fits from a

Michaelis–Menten allometric model in which asymptotic height (*V*m), the scaling exponent (*x*), and the half-saturation diameter (*K*) all vary by infestation outcome; ribbons show 95% credible intervals. Fitted height–DBH trajectories were broadly similar between groups, although trees that subsequently gained infestation showed a weak tendency towards reduced height for a given diameter.

#### Peak infestation intensity *vs.* infestation duration

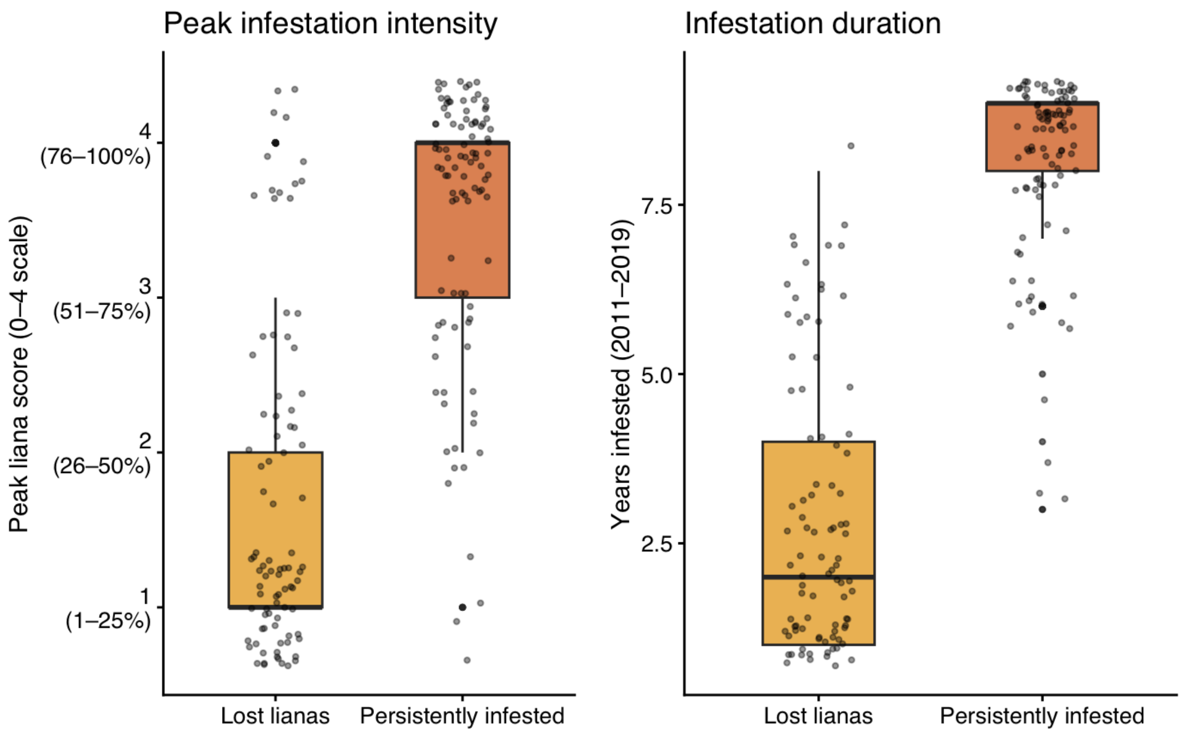


**Figure S2. Trees in the lost-lianas category had lower peak infestation scores and shorter infestation durations than persistently infested trees, suggesting lower cumulative liana burden.** Box plots show medians and interquartile ranges; points show individual trees.

### Random effects (species specific effects)

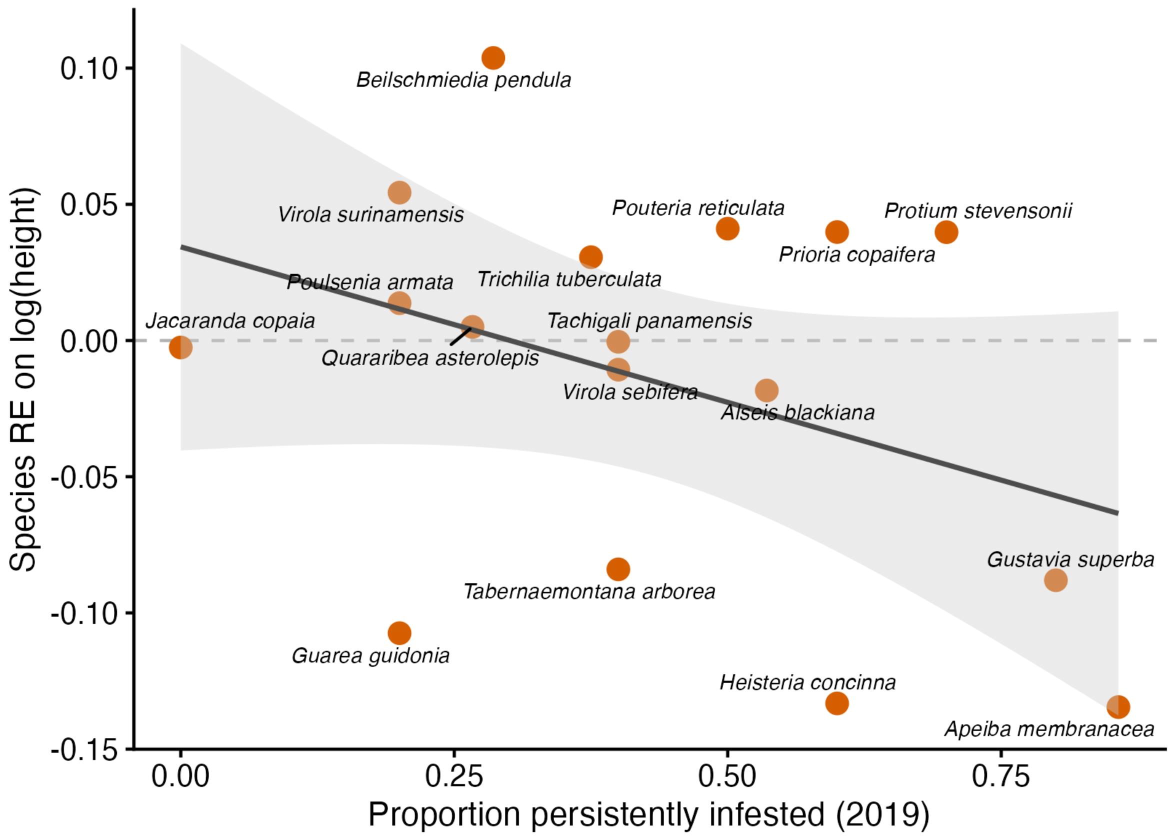


**Figure S3. Exploratory relationship between species random intercepts from the $H_1$ 2019 height model and the proportion of individuals persistently infested.** Points represent species; the fitted line shows the Spearman rank association ($r_s$ = −0.18). The weak negative association suggests that species with higher persistent infestation prevalence tended to have lower height random intercepts after accounting for DBH and infestation history, but uncertainty was high and the pattern should be interpreted cautiously.

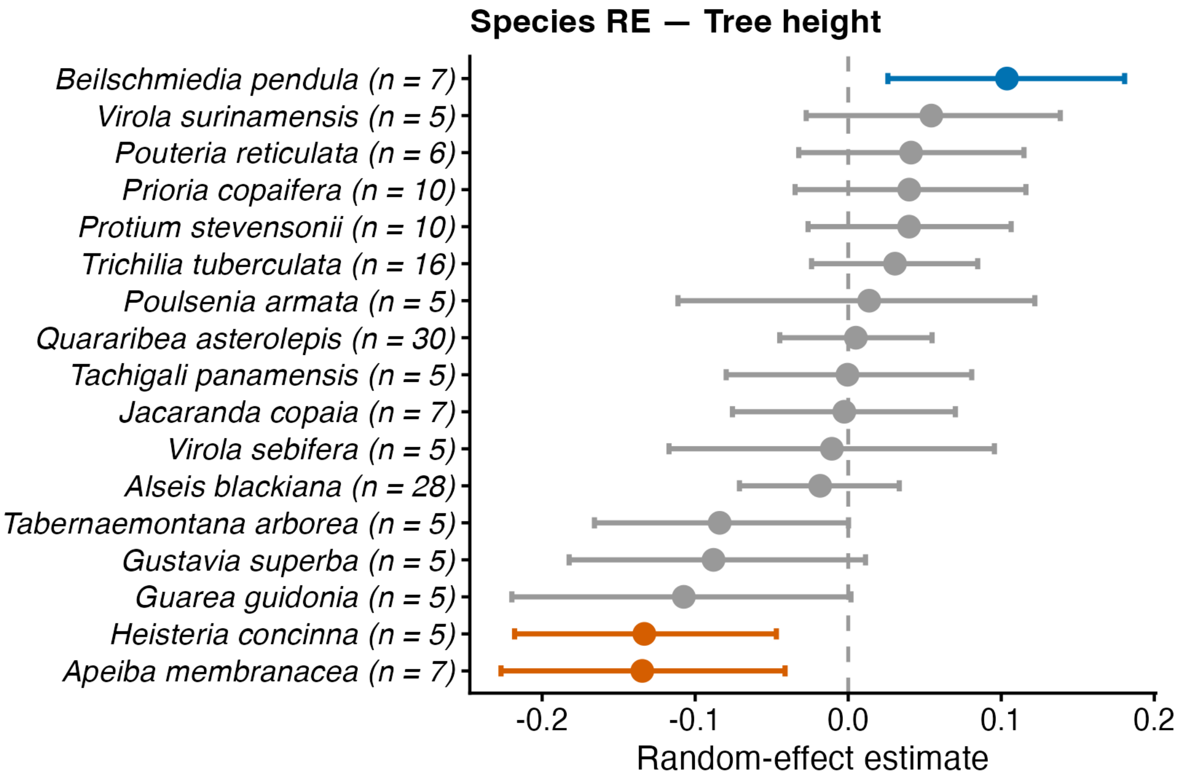


**Figure S4. Species-level variation in tree height after accounting for DBH and infestation history.** Random intercepts from the Bayesian multilevel model for tree height. Points represent species-level deviations from the population mean; horizontal error bars show 95% credible intervals. Blue points indicate species with credibly greater height than the population mean (95% CrI > 0), red points indicate credibly lower height (95% CrI < 0), and grey points indicate no credible deviation from the mean. Estimates are derived from the brms Student-t mixed-effects model including log(DBH), infestation-history category, and random intercepts for species and subplot. Only species with >5 infested individuals and >5 uninfested individuals in 2019 were included.

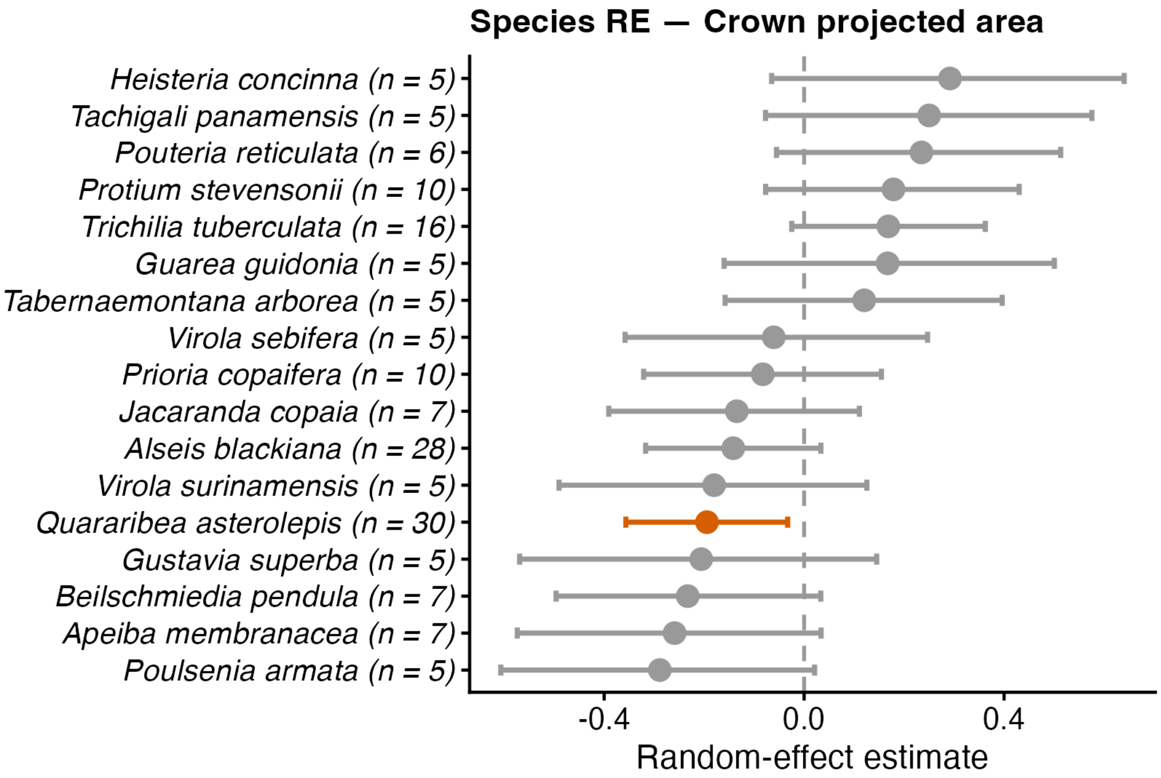


**Figure S5. Species-level variation in crown projected area after accounting for DBH and infestation history.** Random intercepts from the Bayesian multilevel model for crown projected area. Points represent species-level deviations from the population mean; horizontal error bars show 95% credible intervals. Blue points indicate species with credibly greater crown projected area than the population mean (95% CrI > 0), red points indicate credibly lower crown projected area (95% CrI < 0), and grey points indicate no credible deviation from the mean. Estimates are derived from the brms Student-t mixed-effects model including log(DBH), infestation-history category, and random intercepts for species and subplot. Only species with >5 infested individuals and >5 uninfested individuals in 2019 were included.

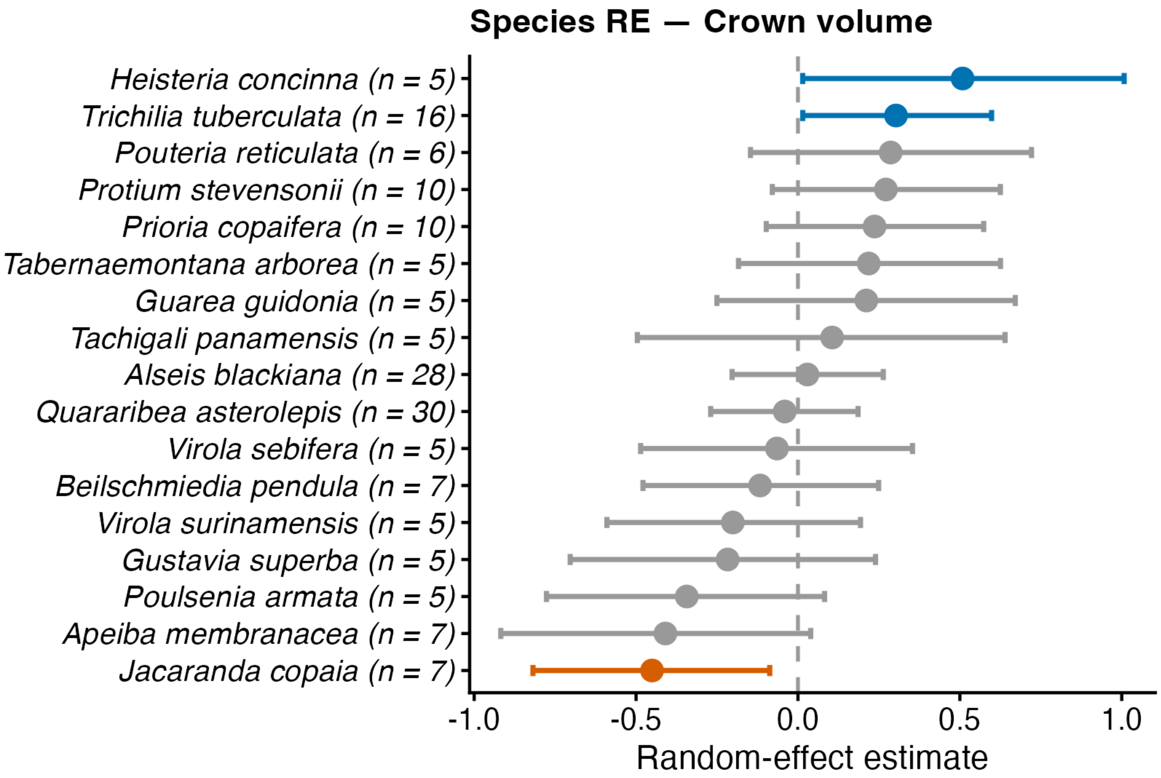


**Figure S6. Species-level variation in crown volume after accounting for DBH and infestation history.** Random intercepts from the Bayesian multilevel model for crown volume. Points represent species-level deviations from the population mean; horizontal error bars show 95% credible intervals. Blue points indicate species with credibly greater crown volume than the population mean (95% CrI > 0), red points indicate credibly lower crown volume (95% CrI < 0), and grey points indicate no credible deviation from the mean. Estimates are derived from the brms Student-t mixed-effects model including log(DBH), infestation-history category, and random intercepts for species and subplot. Only species with >5 infested individuals and >5 uninfested individuals in 2019 were included.

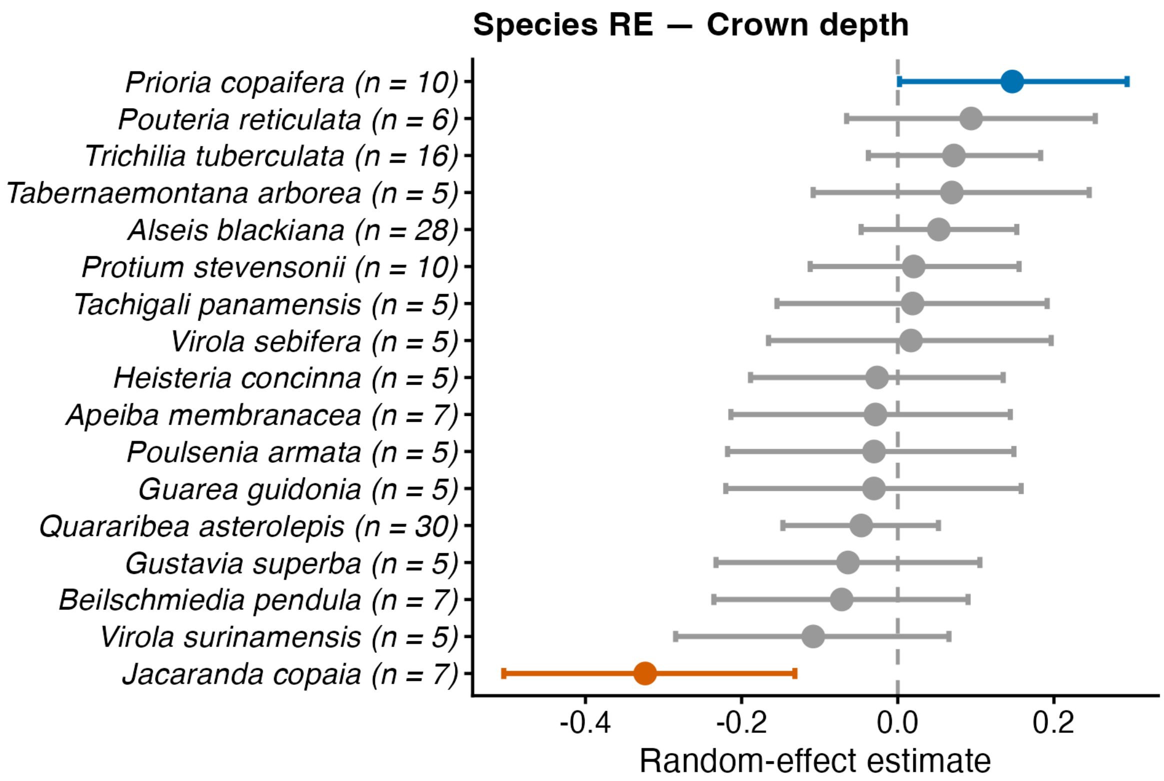


**Figure S7. Species-level variation in crown depth after accounting for DBH and infestation history.** Random intercepts from the Bayesian multilevel model for crown depth. Points represent species-level deviations from the population mean; horizontal error bars show 95% credible intervals. Blue points indicate species with credibly greater crown depth than the population mean (95% CrI > 0), red points indicate credibly lower crown depth (95% CrI < 0), and grey points indicate no credible deviation from the mean. Estimates are derived from the brms Student-t mixed-effects model including log(DBH), infestation-history category, and random intercepts for species and subplot. Only species with >5 infested individuals and >5 uninfested individuals in 2019 were included.

**Climbing modes/ liana infestation pathway**

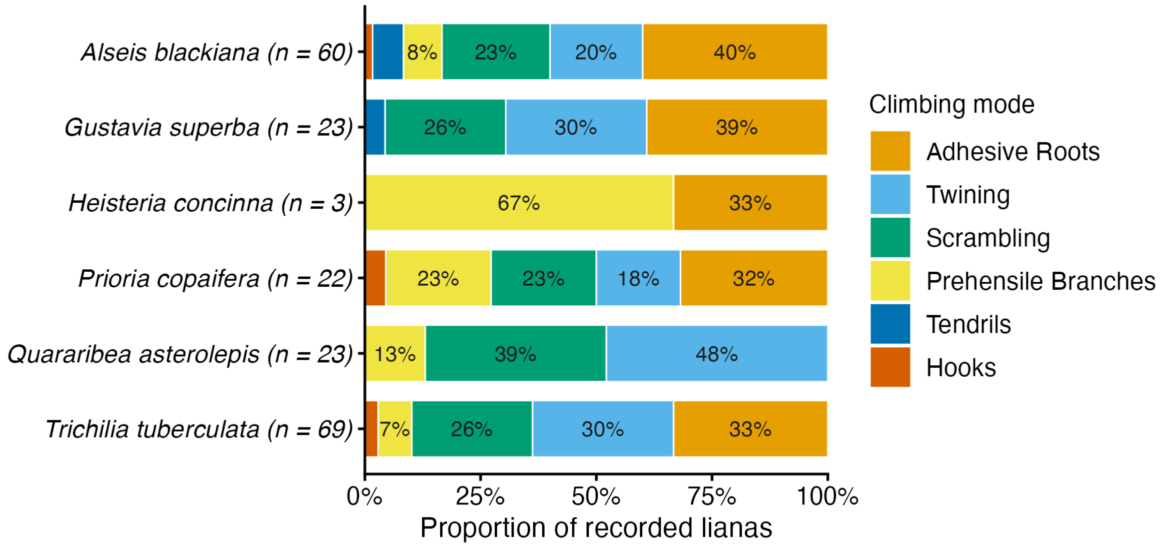


**Figure S8. Climbing modes differed among focal host-tree species surveyed during 2025 field observations.** Proportions show the relative frequency of six liana climbing modes recorded on infested individuals across the focal species dataset. Categories follow Sperotto et al. (2020) and include twiners, tendril climbers, hook climbers, root climbers, adhesive roots, and prehensile branches. Percentages are calculated from all recorded lianas associated with focal host trees during the 2025 survey.

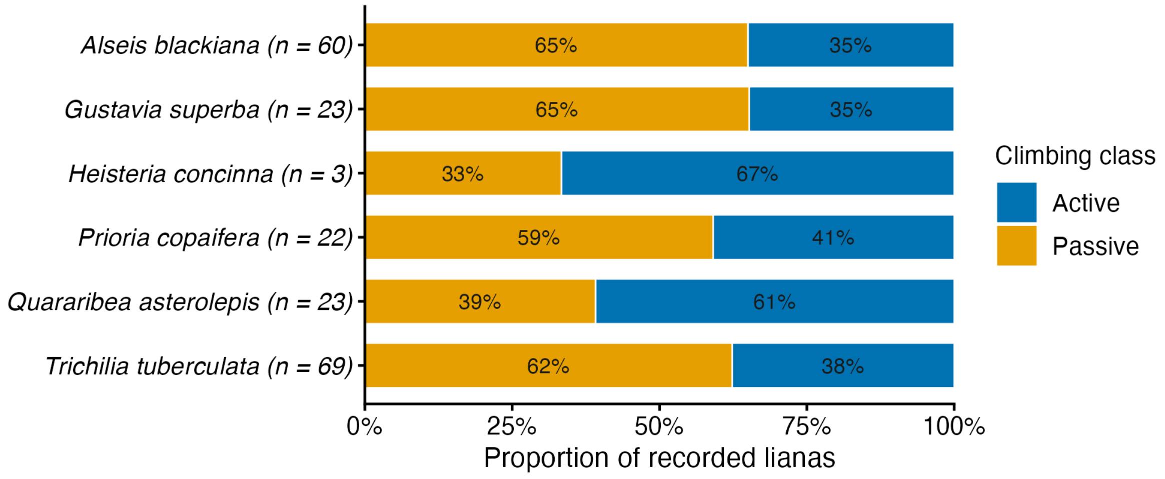


**Figure S9. Active climbing mechanisms dominated liana attachment strategies across focal host-tree species.** Proportions show the frequency of lianas classified as active climbers (twiners, tendril climbers, hook climbers) or passive climbers (adhesive roots, prehensile branches, root climbers) among infested focal trees surveyed in 2025. Percentages are shown separately for each focal host-tree species.

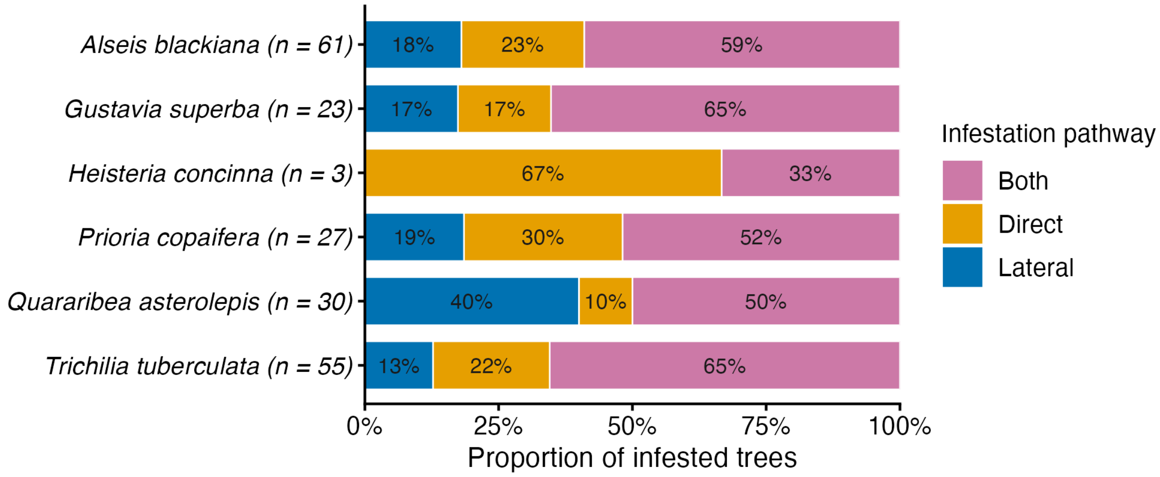


**Figure S10. Most newly infested trees were colonised through both direct ascent and lateral canopy transfer.** Proportions show the infestation pathway recorded for focal host trees during 2025 field surveys. Categories indicate trees colonised through direct ascent from the ground, lateral spread from neighbouring crowns, or both pathways simultaneously. Percentages are shown separately for each focal host-tree species.
